# Identification of novel antiviral drug candidates using an optimized SARS-CoV-2 phenotypic screening platform

**DOI:** 10.1101/2022.07.17.500346

**Authors:** Denisa Bojkova, Philipp Reus, Leona Panosch, Marco Bechtel, Tamara Rothenburger, Joshua Kandler, Annika Pfeiffer, Julian U.G. Wagner, Mariana Shumliakivska, Stefanie Dimmeler, Ruth Olmer, Ulrich Martin, Florian Vondran, Tuna Toptan, Florian Rothweiler, Richard Zehner, Holger Rabenau, Karen L. Osman, Steven T. Pullan, Miles Carroll, Richard Stack, Sandra Ciesek, Mark N Wass, Martin Michaelis, Jindrich Cinatl

**Affiliations:** Institute of Medical Virology, University Hospital, Goethe-University, Frankfurt am Main, Germany; Fraunhofer Institute for Translational Medicine and Pharmacology (ITMP), Frankfurt am Main, Germany; Institute for Cardiovascular Regeneration, Centre of Molecular Medicine, Goethe University, Frankfurt am Main, Germany; German Center for Cardiovascular Research (DZHK), partner site Rhine-Main, Goethe University, Frankfurt am Main, Germany; Cardiopulmonary Institute (CPI), Goethe University, Frankfurt am Main, Germany; Leibniz Research Laboratories for Biotechnology and Artificial Organs (LEBAO), Hannover Medical School, Germany; Clinic for General, Abdominal and Transplant Surgery, Hannover Medical School, Germany; German Center for Infection Research, DZIF, Braunschweig, Germany; Dr Petra Joh Research Institute, Frankfurt am Main, Germany; Institute for Forensic Medicine, University Hospital, Goethe-University, Frankfurt am Main, Germany; Public Health England, National Infection Service, Porton Down, Salisbury, UK; Wellcome Trust Centre for Human Genetics, Nuffield Department of Medicine, Oxford University, Oxford, UK; NIHR Health Protection Unit in Emerging and Zoonotic Infections, Department of Clinical Infection, Microbiology and Immunology, University of Liverpool, Liverpool, UK; School of Biosciences, University of Kent, Canterbury, UK

## Abstract

Reliable, easy-to-handle phenotypic screening platforms are needed for the identification of anti-SARS-CoV-2 compounds. Here, we present caspase 3/7 activity as a read-out for monitoring the replication of SARS-CoV-2 isolates from different variants, including a remdesivir-resistant strain, and of other coronaviruses in a broad range of cell culture models, independently of cytopathogenic effect formation. Compared to other cell culture models, the Caco-2 subline Caco-2-F03 displayed superior performance, as it possesses a stable SARS-CoV-2 susceptible phenotype and does not produce false-positive hits due to drug-induced phospholipidosis. A proof-of-concept screen of 1796 kinase inhibitors identified known and novel antiviral drug candidates including inhibitors of PHGDH, CLK-1, and CSF1R. The activity of the PHGDH inhibitor NCT-503 was further increased in combination with the HK2 inhibitor 2-deoxy-D-glucose, which is in clinical development for COVID-19. In conclusion, caspase 3/7 activity detection in SARS-CoV-2-infected Caco-2F03 cells provides a simple phenotypic high-throughput screening platform for SARS-CoV-2 drug candidates that reduces false positive hits.

## Introduction

There is an ongoing search for antiviral drugs against SARS-CoV-2 that can complement the currently available monoclonal antibody preparations and the three approved small-molecule drugs remdesivir, molnupiravir, and nirmatrelvir [Gao & Sun, 2021]. Effective antiviral drugs and drug combinations will be particularly important for immunocompromised individuals, who cannot effectively be protected by vaccination [Gentile & Schiano Moriello, 2022].

Previous research has shown that the efficacy of antiviral agents may differ between SARS-CoV-2 variants and cell culture models [Dittmar et al., 2021; Bojkova et al., 2022; Zhao et al., 2022]. Some cell culture models may produce false positive hits due to unspecific effects on the host cell metabolism such as phospholipidosis that do not translate into in vivo activity [Tummino et al., 2021]. Moreover, continued SARS-CoV-2 passaging in cell culture may change virus biology, including virus sensitivity to antiviral drugs [Ogando et al., 2020; Ramirez et al., 2021; Szemiel et al., 2021]. Thus, simple and robust cell culture assays that can cover a broad spectrum of SARS-CoV-2 variants (including primary clinical isolates) are required to accelerate the identification of anti-SARS-CoV-2 drug candidates.

Many assays measure SARS-CoV-2-induced host cell destruction (cytopathic effect, CPE) or host cell viability for the identification of antiviral agents [Bojkova et al. 2020; Riva et al., 2020; Touret et al., 2020; Zhang et al., 2020; Ellinger et al., 2021; Van Damme et al., 2021; Yan et al., 2021a]. However, such assays are not suitable for SARS-CoV-2 culture systems that do not display virus-induced cytotoxicity [Caccuri et al., 2020; Liao et al., 2020; Bielarz et al., 2021; Wurtz et al., 2021].

Antibody-based detection of viral antigens and/ or double-stranded RNA is an alternative approach [Dittmar et al., 2021; Garcia et al. 2021], but requires more manual handling. Assays using genetically modified cells, genetically modified SARS-CoV-2 strains, and SARS-CoV-2 replicons have also been developed [Thi Nhu Tao et al., 2020; Xie et al., 2020; He et al., 2021; Van Damme et al., 2021], but cover only the limited number of virus strains that they have been established for.

An ideal assay would enable high throughput screening of wild-type SARS-CoV-2 including the most current clinical isolates in all available cell culture systems in a very simple format that can be applied by many research groups. Such an assay would also enable the phenotypic resistance testing of virus isolates, which is relevant given that the use of antiviral drugs seems to be inevitably associated with the formation of resistant virus variants [Hiscox et al., 2021; Szemiel et al., 2021; Yang et al., 2022].

Here, we introduce an effective screening assay for the identification of compounds that inhibit SARS-CoV-2 replication based on measuring caspase 3/7 activity using a one-step read-out assay (Caspase-Glo® 3/7 Assay System, Promega). This assay works across different coronaviruses including many SARS-CoV-2 strains and clinical isolates as well as across a broad range of cell culture models, including those in which SARS-CoV-2 infection does not result in a recognizable virus CPE. The Caco-2 subline Caco-2-F03 was identified as preferred cell culture model, as it is easy-to-handle, displays a stable susceptibility phenotype, and does not produce false positives due to drug-induced phospholipidosis. A validation screen of 1796 kinase inhibitors confirmed the suitability of our platform and identified 81 compounds that reduced virus-induced caspase activation by more than 90%, including known as well as novel drug candidates such as PHGDH, CLK-1, and CSFR inhibitors.

## Results

### Caco-2 cells as SARS-CoV-2 infection model

The human colon carcinoma Caco-2 cell line was established by Jorgen Fogh (Memorial Sloan-Kettering Cancer Center, New York) in 1974 [Fogh et al., 1977] and has been used for the cultivation of human pathogenic viruses including influenza viruses and coronaviruses since 1985 [Reigel, 1985; Collins, 1990; Chan et al., 2013a]. We already used Caco-2 cells (obtained from DSMZ, Braunschweig, Germany) for the cultivation of the close SARS-CoV-2 relative SARS-CoV starting in 2003 [Cinatl et al., 2003; Cinatl et al., 2004] and they also enabled us and others to quickly cultivate SARS-CoV-2 isolates when this novel virus emerged [Bojkova et al., 2020; Bojkova et al., 2020a; Bojkova et al., 2020b; Hoehl et al., 2020; Klann et al., 2020; Toptan et al., 2020; Bojkova et al., 2021; Ellinger et al., 2021; Gower et al., 2021; Widera et al., 2021].

In our hands, Caco-2 cells (obtained from DSMZ, Braunschweig, Germany at the time) have been highly permissive to SARS-CoV and SARS-CoV-2 and developed a pronounced cytopathogenic effect (CPE) in response to infection with both viruses [Cinatl et al., 2003; Cinatl et al., 2004; Bojkova et al., 2020b; Bojkova et al., 2021]. In other studies, however, Caco-2 cells displayed low SARS-CoV-2 susceptibility and no CPE formation [Chu et al., 2020; Lee et al., 2020; Yeung et al., 2021].

To further investigate these discrepancies, we ordered fresh Caco-2 cells from the following sources: DSMZ (Braunschweig, Germany, designated as Caco-2A), Sigma (Taufkirchen, Germany, Caco-2B), and CLS (Eppelheim, Germany, Caco-2C). To discriminate our original Caco-2 cell line from these other ones, we will refer to it as Caco-2-F03 from now on.

An initial short tandem repeat (STR) analysis confirmed that all Caco-2 cell lines share the reference profile (Suppl. Table 1). However, Caco-2A, Caco2-B, and Caco-2C cells displayed low SARS-CoV-2 permissiveness as indicated by low viral spike (S) protein levels and a lack of CPE formation compared to Caco-2-F03 (Figure 1A, Figure 1B).

**Figure 1.**
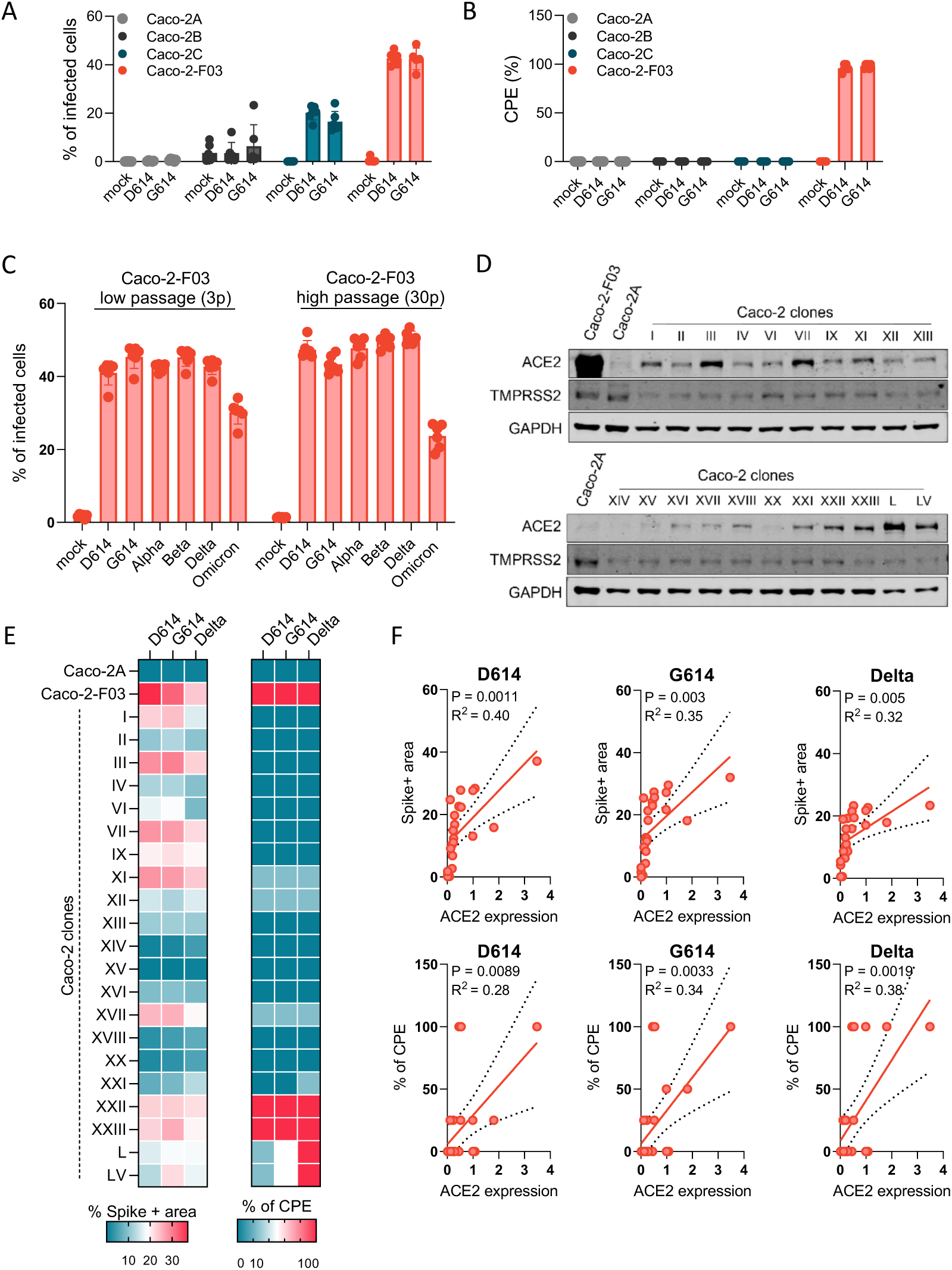
Susceptibility of Caco-2 cells to SARS-CoV-2 infection. A) Percentage of SARS-CoV-2-infected cells detected in Caco-2 cell lines from different sources infected with different SARS-CoV-2 isolates at a multiplicity of infection (MOI) 0.01 as determined by immunostaining for the viral spike (S protein) 48h post infection. B) Cytopathogenic effect (CPE) formation in SARS-CoV-2 (MOI 0.01)-infected Caco-2 cell lines from different sources as determined 48h post infection. C) Susceptibility of Caco-2-F03 cells to a broad range of SARS-CoV-2 isolates after different times of cultivation. Cells had been frozen at passage 14 and were now resuscitated and cultivated for a further 30 passages. SARS-CoV-2 susceptibility was determined by immunostaining for S 48h after SARS-CoV-2 (MOI 0.01) infection 3 and 30 passages post resuscitation. D) ACE2 and TMPRSS2 levels in Caco-2-F03, Caco-2A, and single-cell derived clones from Caco-2A. E) Susceptibility of Caco-2A clones to selected SARS-CoV-2 isolates as indicated by immunostaining for S and CPE formation in SARS-CoV-2 (MOI 0.01)-infected cells 48h post-infection. F) Correlation of S staining and CPE formation with cellular ACE2 levels.

Caco-2-F03 cells remained permissive to SARS-CoV-2 for 30 passages after the resuscitation of cells that had been frozen at passage 14 (Figure 1C), suggesting that their SARS-CoV-2 permissiveness phenotype is stable during prolonged culturing. In agreement, we have used Caco-2-F03 cells since 2003 for the cultivation of initially SARS-CoV and later SARS-CoV-2 [Cinatl et al., 2003; Cinatl et al., 2004; Bojkova et al., 2020b; Bojkova et al., 2021].

Further investigations revealed that Caco-2-F03 cells display high levels of the cellular SARS-CoV and SARS-CoV-2 receptor ACE2 and the protease TMPRSS2, which cleaves and activates S for ACE2 binding [Hoffmann et al., 2020], than Caco-2A, Caco-2B, and Caco-2C (Figure 1D, Suppl. Figure 1).

### The Caco-2A cell line contains SARS-CoV-2-susceptible subpopulations

One explanation for these differences between Caco-2-F03 and Caco-2A, Caco-2B, and Caco-2C is that we may have inadvertently enriched a SARS-CoV-2-permissive subpopulation during Caco-2 cultivation. To test this hypothesis, we established 21 single cell-derived clones from Caco-2A by limited dilution. Four of these clonal sublines were highly susceptible to SARS-CoV-2 infection as demonstrated by high S protein levels and CPE formation (Figure 1E), supporting the hypothesis that Caco-2-F03 has been derived from a SARS-CoV- and SARS-CoV-2-permissive subpopulation of our original Caco-2 cell line. There was some level of correlation between the SARS-CoV-2 susceptibility of Caco-2A clones and the cellular ACE2 levels (Figure 1F) but not between SARS-CoV-2 susceptibility and the cellular TMPRSS2 levels (Suppl. Figure 1). This suggests that ACE2 levels are more important for the SARS-CoV-2 susceptibility of Caco-2 cells than TMPRSS2 levels and that additional mechanisms are also likely to be involved.

### Caspase 3/7 activity for the quantification of the replication of SARS-CoV-2 and other coronaviruses

Coronavirus replication, including that of SARS-CoV-2, has been shown to result in the activation of caspases including the initiator caspases 8 and 9 and the effector caspase 3 [Conolly & Fearnhead, 2017; Bojkova et al. 2020a; Bojkova et al. 2020b; Li et al., 2020; Ren et al., 2020].

Quantification of caspase activity in Caco-2-F03 cells infected with different SARS-CoV-2 isolates at a multiplicity of infection (MOI) of 0.01 48h post infection using the Caspase-Glo assay kit (Promega) resulted in substantially higher signal-to-basal (S/B) ratios for caspase-3/7 (5.9- to 7.7-fold) than for caspase-8 (1.3- to 1.6-fold) and caspase-9 (1.5- to 2.2-fold) (Figure 2A). Caspase 3/7 activity also resulted in higher Z’ scores (0.7-0.9) than caspase 8 (0.5-0.7) and caspase 9 (0.3-0.8) activity (Figure 2A), indicating higher assay robustness [Zhang et al., 1999). Hence, caspase 3/7 activity detection was selected for further investigation as potential screening endpoint method.

**Figure 2.**
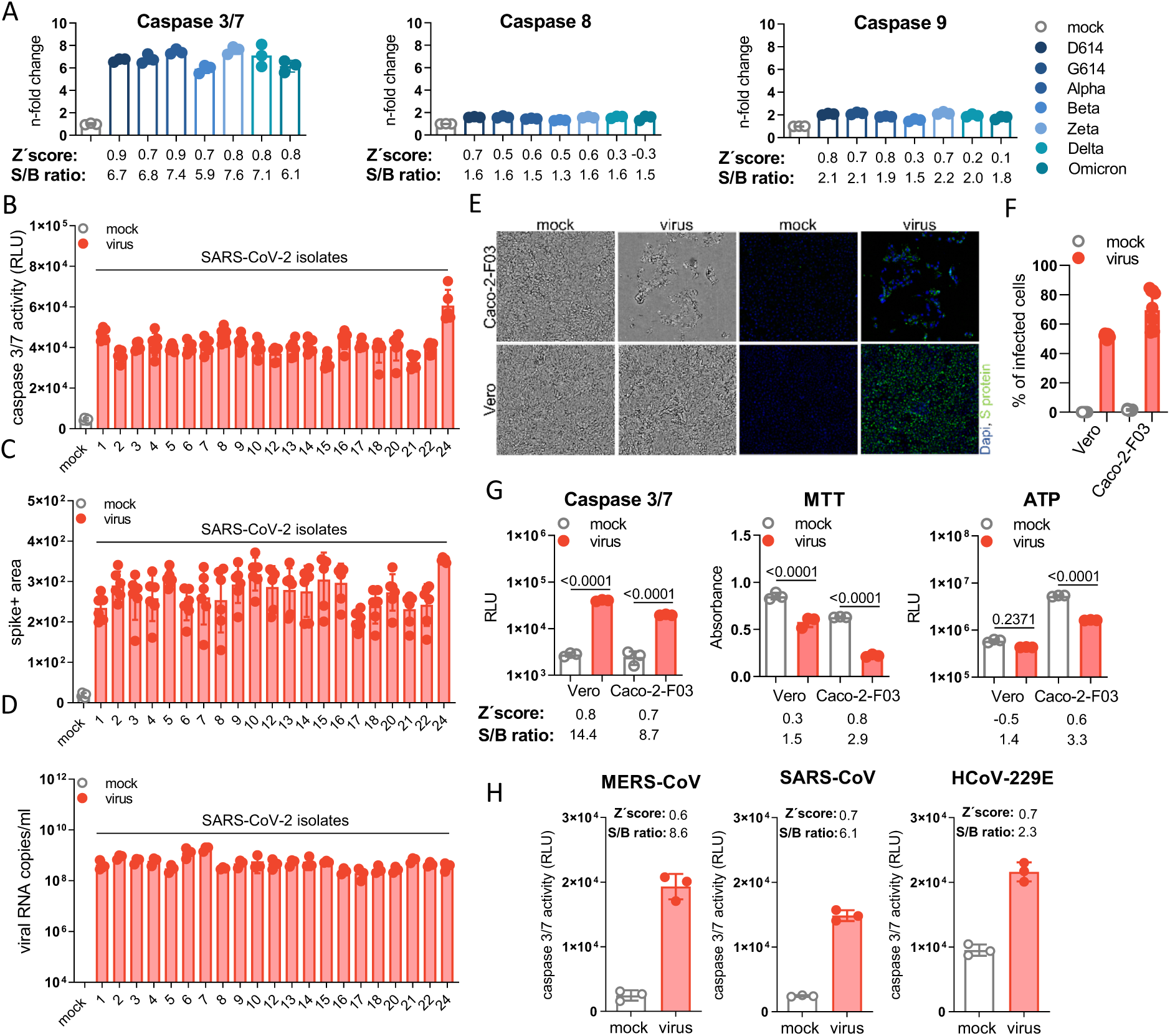
Caspase 3/7 activity for the quantification of the replication of SARS-CoV-2 and other coronaviruses. A) Caspase 3/7, caspase 8, and caspase 9 activity in Caco-2-F03 cells infected with a range of different SARS-CoV-2 isolates (MOI 0.01), as determined by Caspase-Glo assay assay (Promega) 48h post infection. Higher signal-to-basal (S/B) ratios and Z’ scores indicate higher assay robustness. B) Caspase 3/7 activity as determined by Caspase-Glo assay, C) SARS-CoV-2 Spike (S) protein staining, and D) virus titers as indicated by genomic RNA copy numbers determined by qPCR in Caco-2-F03 cells infected with a wide range of uncharacterized SARS-CoV-2 isolates (MOI 0.01) 48h post infection. E) Representative images indicating CPE formation in G614 (MOI 0.01)-infected Caco-2-F03 and Vero cells 48h post infection as indicated by phase contrast microscopy and immunofluoresce staining for the viral S protein in combination with DAPI-stained nuclei. F) Quantification of cellular S protein levels in Caco-2-F03 cells infected with G614 (MOI 0.01) 48h post infection by immunostaining. G) Only caspase 3/7 activity but not viability assays (MTT, CellTiter-Glo measuring ATP production) reflects G614 (MOI 0.01) replication 48h post infection in Vero cells, which do not display a virus-induced CPE. G614 (MOI 0.01)-infected Caco-2-F03 cells served as a control that displays a CPE. P values were calculated by one-way ANOVA. H) Caspase 3/7 activity in Caco-2-F03 cells infected with MERS-CoV, SARS-CoV, and HCoV-229E (MOI 0.01) as determined 48h post infection including S/B ratios and Z’ scores.

Caspase 3/7 activity displayed an MOI-dependent increase in Caco-2-F03 cells 24 h post infection (Suppl. Figure 2A), which mirrored CPE formation (Suppl. Figure 2B). 48h post infection, such differences were not detectable anymore (Suppl. Figure 2A, Suppl. Figure 2B). Moreover, infection of Caco-2-F03 cells with an additional 21 clinical SARS-CoV-2 isolates (after a maximum of two passages in Caco-2-F03 cells) also resulted in effective caspase 3/7 activation (Figure 2B), which reflected viral spike (S) protein levels and virus titers (as indicated by genomic RNA copy numbers) (Figure 2C, Figure 2D). UV-inactivated virus did not cause caspase 3/7 activation (Suppl. Figure 2C), further confirming that SARS-CoV-2-induced caspase 3/7 activation depends on virus replication.

However, caspase 3/7 activation does not appear to be critically involved in SARS-CoV-2 replication as the clinically approved pan-caspase inhibitor emricasan [Shiffman et al., 2010] did not interfere with SARS-CoV-2 replication and CPE formation, despite suppressing caspase 3/7 activity (Suppl. Figure 3).

### Caspase 3/7 activity for the monitoring of SARS-CoV-2 replication in the presence and absence of a virus-induced cytopathogenic effects (CPE)

The caspase 3/7 assay also enabled the monitoring of SARS-CoV-2 replication in Vero cells, in which SARS-CoV-2 does not induce a CPE and in which SARS-CoV-2 replication cannot be monitored by viability assays such as the MTT assay (measures oxidative phosphorylation in the mitochondria) and the Cell TiterGlo assay (Promega, measures cellular ATP production) (Figure 2E-G).

### Caspase 3/7-induction by additional coronaviruses

Caspase 3/7 activity also indicated replication of the additional human-pathogenic coronaviruses SARS-CoV, MERS-CoV, and HCoV-229E in CaCo-2-F03 cells in Caco-2-F03 cells (Figure 2H).

### Caspase 3/7 activity for the monitoring of SARS-CoV-2 replication in primary human cell cultures

Furthermore, SARS-CoV-2-induced caspase 3/7 activity was detected in primary cultures of normal human cells, including induced pluripotent stem cell-derived cardiomyocytes (CMS), air liquid interface (ALI) cultures of bronchial epithelial (HBE) cells, and hepatocytes (PHH) (Suppl. Figure 4A). Immunoblots for the viral nucleoprotein (NP) were used to confirm SARS-CoV-2 infection in these primary cell cultures (Suppl. Figure 4B). ALI HBE did not display disruption of cellular barrier during SARS-CoV-2 infection as measured by transepithelial electrical resistance (TEER) and LDH release (Suppl. Figure 4C and D), whereas CMS and PHH displayed a CPE in response to SARS-CoV-2 infection (Suppl. Figure 4E and F). This is agrees with our previous results that caspase 3/7 activity is a suitable read-out method for monitoring SARS-CoV-2 infection both in the presence and absence of a virus-induced CPE.

Taken together, detection of caspase 3/7 activity does not only enable the monitoring of the replication of a wide range of SARS-CoV-2 variants and clinical isolates (and of other coronaviruses). It is also a suitable read-out for SARS-CoV-2 replication across many different susceptible permanent cell lines and human primary cultures, independently of whether SARS-CoV-2 induces a CPE in these systems.

### Caspase 3/7 activity for the identification of antiviral drugs

Next, we compared caspase 3/7 activity and S protein staining for the detection of the antiviral activity of drugs with known efficacy against SARS-CoV-2, including remdesivir (RNA-dependent RNA polymerase (RdRp) inhibitor), EIDD-1931 (active form of molnupiravir that induces ‘error catastrophe’ in newly produced SARS-CoV-2 genomes), ribavirin (broad-spectrum antiviral drug), nirmatrelvir (3C-like protease/main protease inhibitor), and nafamostat (TMPRSS2 inhibitor) [Apaydın et al., 2021; Simonis et al., 2021] in SARS-CoV-2 variant G614-infected Caco-2-F03 cells. Both detection methods resulted in very similar IC50s (concentrations that inhibit virus activity by 50%) (Figure 3A).

**Figure 3.**
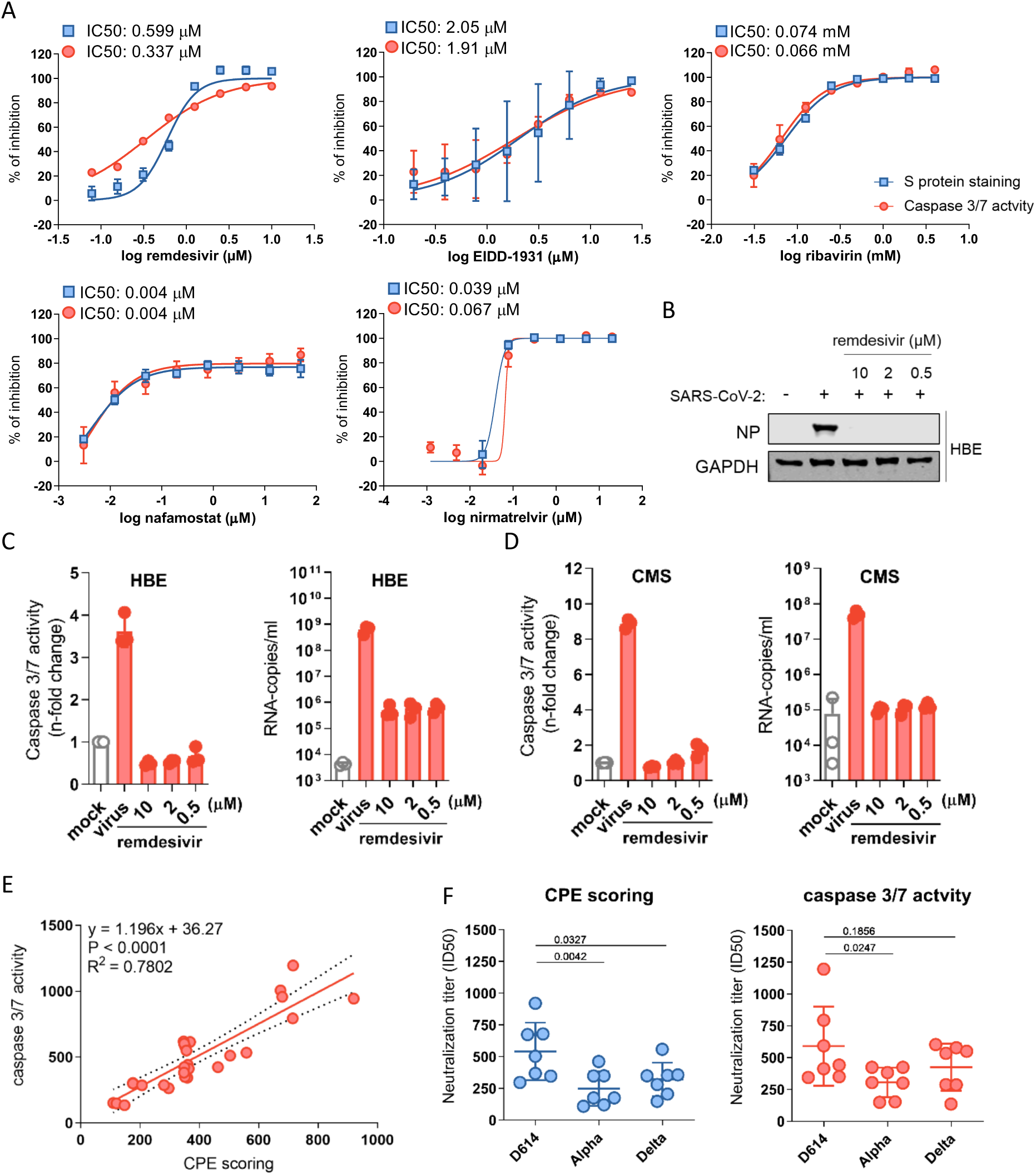
Caspase 3/7 activity for the determination of the antiviral activity of anti-SARS-CoV-2 agents and neutralization assays. A) Dose-response curves and concentrations that inhibit virus infection by 50% (IC50) of antiviral agents as determined by caspase 3/7 activity and immunostaining for the coronavirus S protein in G614 (MOI 0.01)-infected Caco-2-F03 cells 24h post infection. B) Effects of the approved anti-SARS-CoV-2 drug remdesivir on cellular levels of the viral NP protein in G614 (MOI 1)-infected air liquid interface (ALI) cultures of primary human bronchial epithelial (HBE) cells 120h post infection. C) Effects of remdesivir on caspase 3/7 activity and virus titers (genomic RNA copy numbers determined by PCR) in G614 (MOI 1)-infected ALI HBE cultures 120h post infection. D) Effects of remdesivir on caspase 3/7 activity and virus titers in G614 (MOI 1)-infected primary human cardiomyocytes (CMS) 48h post infection. E) Correlation of the neutralization capacity of sera derived from seven donors two weeks after their second dose of the mRNA-1273 vaccine determined by caspase 3/7 activity or cytopathogenic effect (CPE) scoring in D614, Alpha and Delta-infected Caco-2-F03 cells 48h post infection. F) Determination of neutralization titers by caspase 3/7 activity or CPE scoring using sera derived from seven donors two weeks after their second dose of the mRNA-1273 vaccine in Caco-2-F03 cells infected with D614, Alpha, and Delta isolates 72h post infection. P values were calculated using paired t-test.

Caspase 3/7 activity also enabled the monitoring of the antiviral activity of remdesivir in SARS-CoV-2-infected primary ALI HBE and HNE cell cultures that do not display a recognizable CPE. The validity of the results obtained by caspase 3/7 assay was confirmed by determining virus titers and Western Blot analysis of SARS-CoV-2 N protein levels (Figure 3B,C). Finally, caspase 3/7 activity reflected the effect of remdesivir on SARS-CoV-2 replication in CMS (Figure 3D).

Taken together, these findings demonstrate that caspase 3/7 activity enables the monitoring of the antiviral drug response in a broad range of cell culture models.

### Caspase 3/7 activity for the determination of neutralizing antibody titers

Caspase 3/7 activity also enabled the determination of neutralizing antibody titers in sera derived from seven donors two weeks after their second dose of the mRNA-1273 vaccine, as indicated by a close correlation with results obtained by CPE scoring in Caco-2-F03 cells infected with different SARS-CoV-2 variants (Figure 3E, F). The neutralization capacity of the sera was higher against the early SARS-CoV-2 strain FFM3 (G614) than against Alpha (B1.1.7) and Gamma (P.2) variant isolates (Figure 3F), which is in line with the immune evasion properties documented for these variants [Zhang et al., 2021].

### Caspase 3/7 activity for detection of SARS-CoV-2 resistance

Next, we tested whether the caspase 3/7 activity assay may be used for phenotypic screens identifying resistant virus strains. To establish a drug-resistant strain, SARS-CoV-2 strain FFM3 (G614) was passaged in the presence of increasing remdesivir concentrations starting with 0.5 µM (the IC50 concentration) until it could be cultivated in the presence of remdesivir 2µM (FFM3^r^REM). FFM3^r^REM displayed a significantly reduced sensitivity to remdesivir, as indicated by determination of cellular S levels in Caco-2-F03 and Calu-3 cells (Figure 4A) and by caspase 3/7 activity in Caco-2-F03 cells (Figure 4B). Interestingly, this remdesivir-resistant strain displayed increased sensitivity to EIDD-1931 and ribavirin relative to the parental strain (Figure 4A,B).

**Figure 4.**
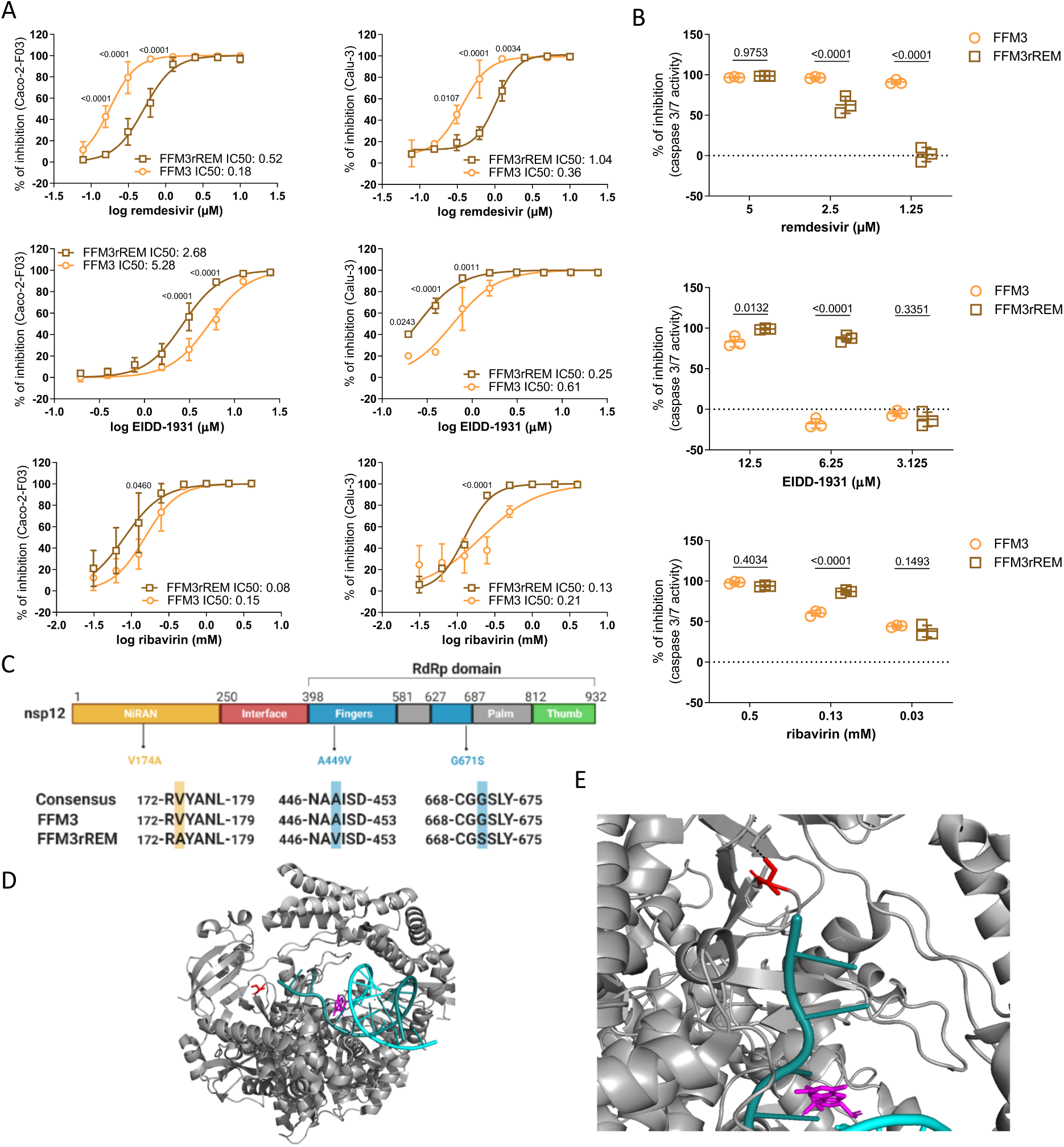
Caspase 3/7 activity for the phenotypic resistance testing of SARS-CoV-2 strains. A) Drug dose response curves and concentrations that reduce cellular levels of the SARS-CoV-2 S protein by 50% (IC50) in Caco-2-F03 and Calu-3 cells as determined by immunostaining 24h (Caco-2-F03) or 48h (Calu-3) post infection with the SARS-CoV-2 strain FFM3 or its remdesivir-adapted substrain FFM3rREM at MOI 0.01. B) Drug concentrations that reduce caspase 3/7 activity in FFM3 and FFM3rREM (MOI 0.01)-infected Caco-2-F03 cells 48h post infection. C) Sequence variants in FFM3rREM compared to FFM3. D) The polymerase complex with nsp7 and nsp8 and a template-primer RNA (cyan and deep teal) and remdesivir (magenta) bound. Gly671Ser is shown in red (as serine). E) The SARS-CoV-2 polymerase Gly671Ser sequence variant. Residue 671 is shown in red as serine, which would be able to form a hydrogen bond with Thr402 which would not be present as Gly671. All p values were calculated by two-way ANOVA.

Sequencing of FFM3^r^REM identified a 154452G>A mutation (present in >90% of alleles) in the coding region of the RNA-dependent RNA polymerase, which results in a change from glycine to serine in position 671(Gly671Ser) (Figure 4C). Gly671Ser is located in the polymerase domain of the RNA-dependent RNA polymerase in close vicinity to where RNA leaves (or enters) the active site (Figure 4D). Gly671Ser could have an effect on the protein structure, as it is located in a bend between two beta sheets, where glycine often has an important role. Additionally, Gly671Ser introduces a side chain capable of forming a hydrogen bond with Thr402 on an adjacent loop (Figure 4E), which could have an effect on the flexibility and conformation of the protein. Therefore, Gly671Ser seems likely to reduce either the binding affinity for remdesivir or to enable the polymerase to overcome the effect of the drug.

Although our structural analysis plausibly explains why Gly671Ser in the RNA-dependent RNA polymerase is likely to mediate remdesivir resistance, it would have been impossible to determine this as a resistance variant without the prior knowledge that the change had happened in response to SARS-CoV-2 adaptation to remdesivir. Hence, this finding emphasizes the relevance of phenotypic assays for the identification of resistant strains that cannot be identified by the analysis of viral genomic information and the subsequent elucidation of the underlying resistance mechanisms. Notably, the caspase 3/7 activity assay also provides an easy-to-use read-out for such phenotypic virus resistance testing approaches.

### Comparison of Caco-2F03 with other cell line candidates for the identification of anti-SARS-CoV-2 drug candidates in screening assays

Next, we directly compared Caco-2F03 to other SARS-CoV-2 cultivation models that could be used for the identification of anti-SARS-CoV-2 drug candidates in screening assays. We focused on permanent cell lines that are easy to cultivate and maintain.

Suitable cell line candidates should be highly permissive for a broad spectrum of SARS-CoV-2 variants and display high caspase 3/7 activity upon infection. We had already shown that Caco-2-F03 cells display high susceptibility to a broad range of SARS-CoV-2 isolates (Figure 2). Here, we directly compared the susceptibility of A549-ACE2, Calu-3, Vero, and Caco-2-F03 cells to D614, G614, Alpha, Beta, and Delta isolates. S immunostaining and caspase 3/7 activity showed that Caco-2-F03 displayed the most pronounced broad-spectrum permissiveness to all tested SARS-CoV-2 isolates (Suppl. Figure 5A, 5B).

Recently, drug-induced phospholipidosis was demonstrated to affect antiviral screens by causing false positive hits due to unspecific effects that do not translate into the clinical setting [Tummino et al., 2021]. In particular, cationic amphiphilic drugs such as hydroxychloroquine were found to induce phospholipidosis [Tummino et al., 2021]. Hence, SARS-CoV-2 culture systems for phenotypic antiviral screening would ideally avoid false positives due to phospholipidosis.

Treatment with hydroxychloroquine resulted in considerable phospholipidosis and inhibited a Delta isolate in A549-ACE2 and Vero cells but not in Calu-3 or CaCo-2-F03 cells (Suppl. Figure 5C, 5D). Due to the susceptibility to the broadest range of SARS-CoV-2 isolates and the insensitivity to drug-induced phospholipidosis, we selected Caco-2-F03 cells for a proof-of-concept screen for anti-SARS-CoV-2 drug candidates.

### Proof-of-concept kinase inhibitor screen for drug candidates that inhibit SARS-CoV-2 replication

Next, we used the caspase 3/7 activity assay in Caco-2F03 cells to screen the Kinase Inhibitor Library (96-well)-L1200 (Selleck) for anti-SARS-CoV-2 drug candidates (Figure 5A, Suppl. File 1). All compounds were tested at a concentration of 10µM. Z’scores were determined as quality controls on all plates as previously described [Xu et al., 2016], and only plates with a Źscore ≥ 0.5 were further analyzed (Figure 5B). Moreover, remdesivir (10 µM) was used as positive control on each plate and produced consistent results (Figure 5B).

**Figure 5.**
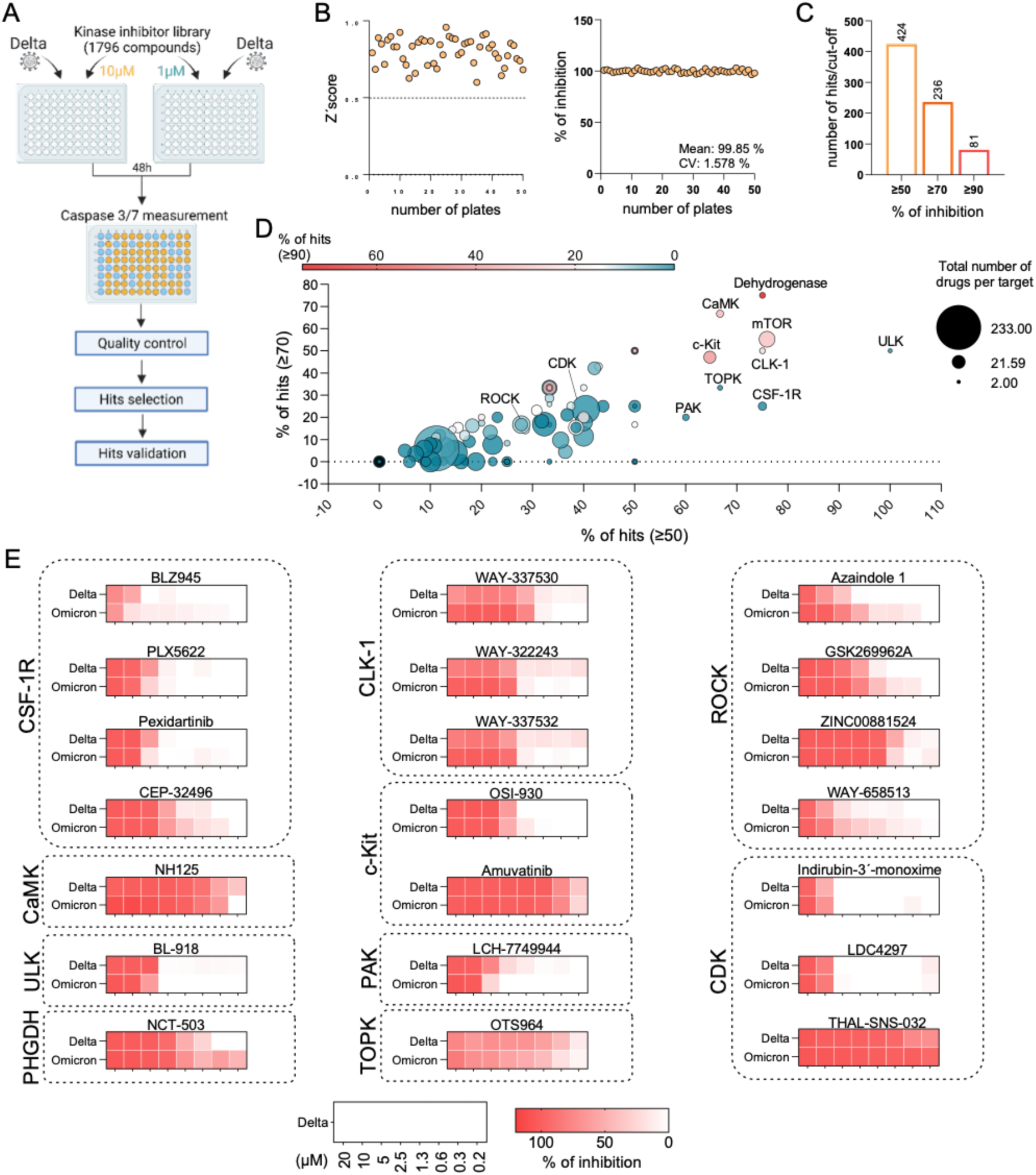
Proof-of-concept screen for anti-SARS-CoV-2 drug candidates using the Caco-2-F03 cell line platform and caspase 3/7 activity as read-out method. A) Overview of the proof-of-concept screen for anti-SARS-CoV-2 compounds using the Kinase inhibitor library L-1200 (Selleckchem, Germany) containing 1796 compounds (Selleckchem, Germany) in Delta (MOI 0.01)-infected Caco-2-F03 cells using caspase 3/7 activity as read-out 48h post infection. For the screen, every compound was tested at a concentration of 10 and 1 µM. 21 selected hits were then confirmed by determining drug-response curves. B) Quality controls, Z’scores served as quality controls (left). Only plates with a Źscore ≥ 0.5 were further analyzed. Remdesivir (10 µM) was used as positive control on each plate and produced consistent results (right). C) Number of hits at different inhibition cut-offs. D) Visualization of the distribution of hits according to their targets. Targets for which inhibitors were selected for confirmation are indicated. E) Heatmaps of the anti-SARS-CoV-2 activity of 21 hits by the determination of dose-response in Delta and Omicron (MOI 0.01)-infected Caco-2-F03 cells using immunostaining for the viral S protein as read-out 24h post infection.

All compounds were tested at a concentration of 10µM, which resulted in 81 hits, when we considered ≥ 90% inhibition of caspase 3/7 activity as a cut-off (Figure 5C). Most hits were identified among inhibitors that target dehydrogenases, CaMK, mTOR, ULK, CLK-1, TOPK, CSF-1R, and PAK (Figure 5D). CaMK, mTOR, ULK, TOPK (also known as PBK), and PAK had already been proposed as antiviral drug targets for SARS-CoV-2 [Shahinozzaman et al., 2020; Jamaly et al., 2021; Shang et al., 2021; Agrawal et al., 2022; Basile et al., 2022]. However, we could not find any information on potential anti-SARS-CoV-2 effects caused by CLK-1 or CSF-1R inhibition. We also included the phosphoglycerate dehydrogenase (PHGDH) inhibitor NCT-503 [Pacold et al., 2016; Hamanaka et al., 2018] in our confirmation experiments. Although dehydrogenases had been known to contribute to SARS-CoV-2 replication [Shang et al., 2021], PHGDH had not previously been shown to be involved.

In addition to inhibitors of the targets described above, ROCK and CDK inhibitors were also included into the confirmation experiments. ROCK was described to be involved in the SARS-CoV-2-induced suppression of the host cell interferon response [Zhang et al., 2021a]. CDK inhibitors had previously been shown to inhibit SARS-CoV-2 replication [Gutierrez-Chamorro et al., 2021; Hahn et al., 2021]. The determination of dose-response curves for all of the 21 inhibitors using immunostaining for the SARS-CoV-2 S protein confirmed the results of the screen (Figure 5E, Suppl. Figure 6).

### PHGDH inhibitor NCT-503 as anti-SARS-CoV-2 drug candidate

Since PHGDH is a new potential antiviral drug target for the treatment of SARS-CoV-2 infection, we further investigated NCT-503. To investigate whether PHGDH inhibition is critical for NCT-503-mediated SARS-CoV-2 inhibition, we compared its effects and those of a chemically closely related analogue, which does not inhibit PHGDH and is commonly used as inactive NCT-503 control (Suppl. Figure 7A) [Pacold et al., 2016; Arlt et al., 2021], for antiviral activity. Only NCT-503 but not the inactive control inhibited Delta- and Omicron-induced caspase 3/7 activation indicating that the antiviral effects of NCT-503 are indeed mediated by PHGDH inhibition (Suppl. Figure 7B).

NCT-503 also inhibited SARS-CoV-2 replication in primary human bronchial epithelial cell air-liquid interface (ALI) cultures (Figure 6A) as indicated by SARS-CoV-2-induced caspase 3 activity (Figure 6B), viral titers (determined as copy numbers of genomic RNA by PCR) (Figure 6C), cell layer integrity (Figure 6D), and lack of SARS-CoV-2-induced cytotoxicity (as indicated by LDH release) (Figure 6E).

**Figure 6.**
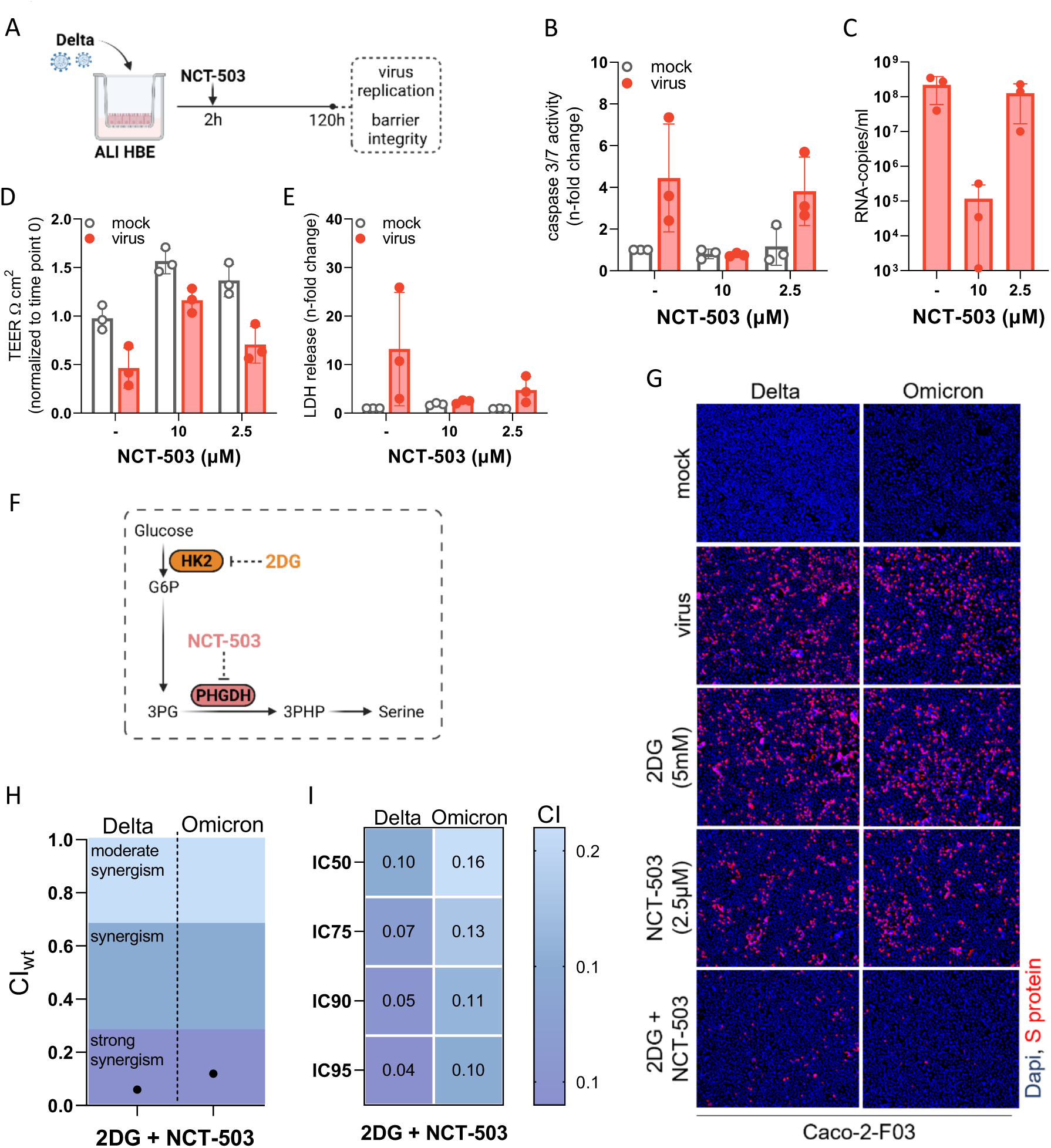
Investigation of the anti-SARS-CoV-2 effects of the PHGDH inhibitor NCT-503 alone or in in combination with 2-Deoxy-D-glucose (2DG). A) Scheme of the testing of NCT-503 for anti-SARS-CoV-2 activity in air liquid interface (ALI) cultures of primary human bronchial epithelial (HBE) cells. Effect of NCT-503 on (B) caspase 3/7 activity, (C) virus titers (determined as genomic RNA copy numbers by qPCR), (D) transepithelial electrical resistance (TEER), and (E) LDH release in ALI HBE cultures infected with Delta (MOI 1) 120h post infection. F) Anti-SARS-CoV-2 effects of NCT-503 in combination with 2-Deoxy-D-glucose (2DG). Illustration of how NCT-503 and 2DG can exert combined effects on a common metabolic pathway. G) Representative fluorescence images indicating the number of Delta and Omicron (MOI 0.01)-infected cells in NCT503 and/ or 2DG-treated Caco-2-F03 cultures 24h post infection. H) and I) Weighted combination indices (CIwt) determined by the method of Chou and Talalay [Chou, 2006] indicating a strong synergism of NCT-503 and 2DG.

Taken together, NCT-503 is a novel antiviral drug candidate for the treatment of SARS-CoV-2 infections that inhibits virus replication via PHGDH inhibition and is effective in different model systems including primary human bronchial epithelial cell ALI cultures, the system considered to be most physiologically relevant [Mulay et al., 2021].

### NCT-503 in combination with 2-Deoxy-D-glucose (2DG)

The discovery of PHGDH as novel antiviral drug target and of NCT-503 as antiviral drug candidate offers potential additional opportunities for combination therapies that display higher efficacy than either single treatment.

*De novo* serine synthesis is a side branch of glycolysis that includes the conversion of the glycolytic intermediate 3-phosphoglycerate (3PG) into 3-phosphohydroxypyruvate (3PHP) by PHGDH (Figure 6F) [Geeraerts et al., 2021]. The production of 3PG in the glycolytic cycle depends on the phosphorylation of glucose into glucose-6-phosphate (G6P) by hexokinase II (HK2) as an initial step (Figure 6F) [Pajak et al., 2020]. Notably, the HK2 inhibitor 2-Deoxy-D-glucose (2DG) has already been shown to inhibit SARS-CoV-2 replication [Bojkova et al., 2020; Bojkova et al., 2021a]. Hence, we hypothesized that the combined inhibition of *de novo* serine synthesis by 2DG and NCT-503 may result in further enhanced antiviral effects (Figure 6F).

Indeed, the combination of 2DG and NCT-503 resulted in stronger Delta and Omicron BA.1 inhibition than either drug alone (Figure 6G, Suppl. Figure 7C). The determination of combination indices (CIs) by the method of Chou and Talalay [Chou, 2006] indicated a strong synergism of NCT-503 and 2DG against both SARS-CoV-2 isolates (Figure 6H, Figure 6I).

## Discussion

Here, we developed a novel screening assay for the identification of anti-SARS-CoV-2 compounds, based on using caspase 3/7 activity determined by the Caspase-Glo^®^ Assay System as read-out indicating SARS-CoV-2 replication. This one step read-out assay can be used by the large number of laboratories, which are equipped with the required plate readers that are in common use. Moreover, the assay is widely (potentially universally) applicable to different SARS-CoV-2 strains and clinical isolates as well as cell culture systems, as indicated by our wide range of pilot experiments. Moreover, our findings show that caspase 3/7 activity can also be used to determine SARS-CoV-2 replication in neutralization assays determining the antibody response in the plasma of individuals and for the phenotypic resistance testing of virus variants. Notably, the caspase 3/7 assay also detects SARS-CoV-2 replication in cultivation systems that do not develop a CPE and in which viability assays such as the MTT assay and the Cell Titer Glo^®^ Assay did not reflect SARS-CoV-2 replication.

For our proof-of-concept experiment for phenotypic resistance testing, we established a remdesivir-resistant SARS-CoV-2 strain by adapting the SARS-CoV-2 strain FFM3 to replication in the presence of remdesivir. Our results confirmed previous observations [Szemiel et al., 2021; Yang et al., 2022] demonstrating that SARS-CoV-2 resistance formation against clinically approved antiviral drugs poses a relevant risk. Notably, the genetic sequence of our remdesivir-adapted SARS-CoV-2 strain would not have enabled us to identify this as a resistant strain by a genotypic approach, which emphasizes the potential need for effective phenotypic resistance testing platforms in the future.

In addition to identifying an easy-to-handle read-out assay for anti-SARS-CoV-2 agent screens, we were also interested in identifying a well-suited cell culture platform. We considered permanent cell lines to be the most promising candidates, because they are readily available and require a minimum of handling. A number of continuous cell lines (e.g. A549-ACE, Calu-3, Vero, Caco-2) had already been used in different phenotypic screening approaches for the identification of antiviral drug candidates against SARS-CoV-2 [Dittmar et al., 2021; Ellinger et al., 2021; Xu et al., 2021]. Based on our comparison of different candidate cell lines, however, we identified Caco-2-F03 as the best platform, as it displayed susceptibility to the widest range of SARS-CoV-2 strains and isolates and was not affected by drug-induced phospholipidosis that has been shown to result in false-positive hits during the testing of anti-SARS-CoV-2 drug candidates [Tummino et al., 2021].

Notably, Caco-2 cells were (in contrast to Calu-3, A549, or Vero cells) shown to be highly susceptible to seasonal coronaviruses such as HCoV-229E or HCoV-OC43 [Collins, 1990; Tang et al., 2005; Yoshikawa et al., 2010; Michaelis et al., 2011; Chan et al., 2013; Ramani et al., 2021]. In this context, we found here that the caspase 3/7 assay also enabled the monitoring of HCoV-229E replication in Caco-2-F03 cells, indicating that this may also serve as a unique broad-spectrum drug screening platform for (seasonal) coronaviruses.

Our study also provided an explanation for the contradictory findings on the SARS-CoV-2 susceptibility of Caco-2 cells reported in previous studies [Bojkova et al., 2020; Bojkova et al., 2020b; Chu et al., 2020; Hoehl et al., 2020; Klann et al., 2020; Lee et al., 2020; Toptan et al., 2020; Bojkova et al., 2021; Ellinger et al., 2021; Gower et al., 2021; Widera et al., 2021; Yeung et al., 2021]. When we investigated newly acquired Caco-2 cell lines from different providers (DSMZ, CLS, Sigma) for SARS-CoV-2 susceptibility, they did indeed not present the level of SARS-CoV-2 permissiveness that we find in our Caco-2-F03 cell line. The subsequent analysis of 21 clonal sublines of the newly purchased lowly SARS-CoV-2-susceptible Caco-2 cell line from DSMZ (Caco-2A) resulted in a broad range of susceptibility phenotypes, suggesting that a highly SARS-CoV-2-susceptible subpopulation has inadvertently become the dominant population in our Caco-2-F03 cell line. Notably, the susceptibility phenotype of Caco-2-F03 appears to be stable, as we have used this cell line for the cultivation of SARS-CoV and SARS-CoV-2 since 2003 [Cinatl et al., 2003; Cinatl et al., 2004]. Moreover, the SARS-CoV-2 susceptibility phenotype of Caco-2-F03 was maintained for a further 30 passages within the current study. Notably, such phenotypic differences between samples of the same cell line obtained from different sources is not surprising and has been described for different cell lines [Feichtinger et al., 2016; Ben-David et al., 2018; Liu et al., 2019].

Next, we used the caspase 3/7 activity assay in Caco-2-F03 cells to screen the Kinase Inhibitor Library (96-well)-L1200 (Selleck) for anti-SARS-CoV-2 drug candidates, which resulted in 81 hits that reduced SARS-CoV-2-induced caspase 3/7 activity by ≥ 90%. These hits included inhibitors of known potential anti-SARS-CoV-2 drug targets (CaMK, mTOR, ULK, TOPK, PAK, ROCK, CDK) [Shahinozzaman et al., 2020; Jamaly et al., 2021; Ellinger et al., 2021; Shang et al., 2021; Zhang et al., 2021b; Agrawal et al., 2022; Basile et al., 2022] and those that interfere with drug targets that had not previously been identified to be relevant during SARS-CoV-2 replication (CLK-1, CSF-1R). We determined dose response curves for 21 out of these 81 hit compounds using immunostaining for the viral S protein, which confirmed their anti-SARS-CoV-2 activities.

Among these hits, we further investigated the phosphoglycerate dehydrogenase (PHGDH) inhibitor NCT-503 [Pacold et al., 2016; Hamanaka et al., 2018], as it interferes with a dehydrogenase that had not previously been shown to be involved in SARS-CoV-2 replication. In addition to NCT-503, we tested a structurally closely related analogue that does not inhibit PHGDH [Pacold et al., 2016; Arlt et al., 2021]. This inactive NCT-503 analogue did not affect SARS-CoV-2 replication, indicating that the anti-SARS-CoV-2 effects of NCT-503 are caused by its effect on PHGDH.

PHGDH activity is critically involved in *de novo* serine synthesis [Geeraerts et al., 2021], a pathway downstream of the glycolytic cycle that depends on the phosphorylation of glucose into glucose-6-phosphate (G6P) by hexokinase II (HK2) as initial step [Pajak et al., 2020]. Since the HK2 inhibitor 2-Deoxy-D-glucose (2DG) has already been shown to inhibit SARS-CoV-2 replication [Bojkova et al., 2020; Bojkova et al., 2021a], we tested whether the combined interference with this pathway using NCT-503 and 2DG resulted in further increased antiviral effects. Indeed, the combination resulted in strongly synergistic anti-SARS-CoV-2 activity. Such antiviral combination therapies have been suggested to be of critical importance for the sustained control of virus outbreaks, as they are not only more effective but also anticipated to reduce and, ideally, prevent resistance formation [White et al., 2021].

In conclusion, we here present a novel phenotypic screening platform for the identification of drug candidates with activity against SARS-CoV-2 and other coronaviruses based on the determination of caspase 3/7 activity using the one-step Caspase-Glo^®^ 3/7 Assay System as read-out. Caspase 3/7 activity is also a suitable read-out for neutralization assays and phenotypic resistance testing. The Caco-2-F03 cell line was identified as the best-suited cell culture platform. It is susceptible to a particularly broad range of SARS-CoV-2 isolates and its susceptibility phenotype remains stable over many passages. Moreover, Caco-2-F03 is not affected by phospholipidosis, which is known to cause false-positive hits during the testing of potential anti-SARS-CoV-2 agents [Tummino et al., 2021]. Hence, the determination of caspase 3/7 activity in SARS-CoV-2-infected Caco-2-F03 cells represents a newly established screening platform that is easy-to-use also for groups without experience in drug discovery projects. A proof-of-concept screen of a kinase inhibitor library containing 1796 compounds resulted in known and novel anti-SARS-CoV-2 drug targets. The PHGDH inhibitor NCT-503 was identified as novel antiviral drug candidate, whose activity was further increased by 2DG (an inhibitor of the PHGDH upstream HK2), which is under clinical development for the treatment of COVID-19 treatment [Sahu & Kumar, 2021].

## Material and methods

### Cell culture

Caco-2A (DSMZ), Caco-2B (Sigma), Caco-2C (CLS), Vero (DSMZ), Calu-3 (ATCC), and Caco-2-F03 (Resistant Cancer Cell Line collection, https://research.kent.ac.uk/industrial-biotechnology-centre/the-resistant-cancer-cell-line-rccl-collection/) were grown at 37 °C in minimal essential medium (MEM) supplemented with 10% fetal bovine serum (FBS), 100 IU/mL penicillin, and 100 μg/mL streptomycin. All culture reagents were purchased from Sigma. A549-ACE2 (Invivogen) was grown in DMEM supplemented with 10% FBS, 2% L-glutamine, 100 μg/ml normocin, 0.5 μg/ml puromycin, 100 IU/mL penicillin, and 100 μg/mL of streptomycin. All cell lines were regularly authenticated by short tandem repeat (STR) analysis and tested for mycoplasma contamination.

Primary bronchial epithelial cells were isolated from the lung explant tissue of a patient with lung emphysema as previously described [van Wetering et al., 2000]. For differentiation into air-liquid interface (ALI) cultures, cells were resuscitated, passaged once in PneumaCult-Ex Medium (StemCell technologies), and seeded on transwell inserts (12-well plate, Sarstedt) at 4×10^4^ cells/insert. After reaching confluence, medium on the apical side of the transwell insert was removed and medium in the basal chamber was replaced with PneumaCult ALI Maintenance Medium (StemCell Technologies) including Antibiotic/Antimycotic solution (Sigma Aldrich) and MycoZap Plus PR (Lonza). Criteria for successful differentiation were the development of ciliary movement, an increase in transepithelial electric resistance, and mucus production.

Human induced pluripotent stem cell-derived cardiomyocytes (hiPS-CMs) of two donors were obtained with an embryoid body-based protocol as previously described [Breckwoldt et al., 2017]. hiPS-CMs were cultured in RPMI/B27 medium at 37 °C and 5 % CO2 for 4 to 5 days prior to viral infection.

Primary human hepatocytes (PHHs) were isolated as previously described [Vondran et al., 2008] and were maintained in William’s Medium E (PAN Biotech, Aidenbach, Germany) containing 10% fetal bovine serum (Biochrom, Cambridge, UK) and 10,000 U penicillin/streptomycin, 1% L-glutamine, 1% non-essential amino-acids, 5mmol/L Hepes (Thermo Fisher Scientific, Schwerte, Germany), 2% dimethyl sulfoxide (DMSO, Roth, Karlsruhe, Germany), 5 µg/mL insulin, and 0.05 mmol/L hydrocortisone (Sigma Aldrich, Munich, Germany).

### Virus preparation and infection of different cell types

Caco-2-F03 cells were used for the isolation SARS-CoV-2 variants applied in this study. Information on the following isolates is available from GenBank: D614 (SARS-CoV-2/FFM1, MT358638), G614 (SARS-CoV-2/FFM7, MT358643), Alpha (SARS-CoV-2/FFM-UK7931/2021, MZ427280), Beta (SARS-CoV-2/FFM-ZAF1/2021, MW822592), Delta (SARS-CoV-2/FFM-IND8424/2021, MZ315141), Zeta (SARS-CoV-2/FFMBRA1/2021, MW822593), Omicron (SARS-CoV-2/FFM-SIM0550/2021, OL800702). Additional isolates were not further characterized. SARS-CoV-2 stocks were cultivated for a maximum of three passages in Caco-2-F03 cells and stored at – 80°C. SARS-CoV stocks were prepared on Caco-2-F03 cells as previously described [Cinatl et al., 2004]. MERS-CoV was obtained from BEI Resources (EMC/2012, NR-44260) and passaged once on Vero cells prior experiments. Viral stocks of HCoV-229E (ATCC no. CCL-137) were prepared using Caco-2-F03 cells. Virus titers were determined as TCID50/mL in confluent cells in 96-well microtiter plates.

Primary bronchial and nasal epithelial cells in ALI cultures were infected with SARS-CoV-2 from the apical site. The inoculum was incubated for 2 h, then removed and cells were washed three times with PBS. For testing of antiviral activity of drugs, the compounds were added after the infection period from both the apical and the basal site. The apical medium was removed after one day.

### Caspase activity assay

Caspase 3/7, 8 and 9 activity was measured using the Caspase-Glo assay kit (Promega, Madison, WI, USA), according to the manufacturer’s instructions. Briefly, 100 μL of Caspase-Glo reagent were added to each well, mixed, and incubated at room temperature for 30 min. Luminescence intensity was measured using an Infinite M200 microplate reader (Tecan).

### Viability assay

Cell viability was measured by 3-(4,5-dimethylthiazol-2-yl)-2,5-diphenyltetrazolium bromide (MTT) dye reduction assay. 25 µL of MTT solution (2 mg/mL in PBS) were added per well, and the plates were incubated at 37 °C for 4 h. After this, the cells were lysed using 100 µL of a buffer containing 20% sodium dodecylsulfate and 50% *N*,*N*-dimethylformamide with the pH adjusted to 4.7 at 37 °C for 4 h. Absorbance was determined at 560 nm (reference wavelength 620 nm) using a Tecan infinite M200 microplate reader (TECAN).

Alternatively, cell viability was determined using the CellTiter-Glo (Promega), which measures ATP production, according to the manufacturer’s protocol. Luminescence was measured on a Tecan infinite M200 microplate reader (TECAN).

### Immunocytochemistry of viral antigen

Cells were fixed with acetone:methanol (40:60) solution and immunostaining was performed using a monoclonal antibody directed against the spike (S) protein of SARS-CoV-2 (1:1500, Sinobiological), which was detected with a peroxidase-conjugated anti-rabbit secondary antibody (1:1,000, Dianova), followed by addition of AEC substrate. The S positive area was scanned and quantified by the Bioreader® 7000-F-Z-I microplate reader (Biosys). The results are expressed as percentage of inhibition relative to virus control which received no drug.

### Immunofluorescence labeling

Cells were fixed with 3% PFA permeabilized with 0.1 % Triton X-100. Prior to primary antibody labeling, cells were blocked with 5% donkey serum in PBS or 1% BSA and 2% goat serum in PBS for 30 minutes at room temperature. Spike (S) protein was detected using a specific antibody (1:1500, Sinobiological) and an Alexa Fluor 488 anti-rabbit secondary antibody (1:200, Invitrogen). The nucleus was labeled using DAPI (1:1000, Thermo Scientific). Cardiomyocytes were counterstained with Alexa FluorTM 647 Phalloidin (1:100, #A22287, Invitrogen). Images were taken using Spark® Mulitmode microplate reader (TECAN) at 10x magnification.

### Immunoblot assay

Cells were lysed using Triton-X-100 sample buffer (Sigma-Aldrich), and proteins were separated by SDS-PAGE. Detection occurred by using specific antibodies against GAPDH (1:1000 dilution, #2275-PC-100, Trevigen), SARS-CoV-2 NP (1:1000 dilution, #40143-R019, Sino Biological), ACE2 (1:500 dilution, #ab15348, Abcam), and TMPRSS2 (1:1000 dilution, Recombinant Anti-TMPRSS2 antibody [EPR3861], #ab92323, Abcam) followed by incubation with IRDye-labeled secondary antibodies (LI-COR Biotechnology, IRDye®800CW Goat anti-Rabbit, 926-32211, 1:40,000) according to the manufacturer’s instructions. Protein bands were visualized by laser-induced fluorescence using an infrared scanner for protein quantification (Odyssey, Li-Cor Biosciences, Bad Homburg, Germany).

### qRT-PCR

SARS-CoV-2 RNA from cell culture supernatant samples was isolated using AVL buffer and the QIAamp Viral RNA Kit (QIAGEN) according to the manufacturer’s instructions. Quantification of viral RNA was performed as previously described [Bojkova et al., 2020; Toptan et al., 2020] using primers targeting the RNA-dependent RNA polymerase (RdRp): RdRP_SARSr-F2 (GTG ARA TGG TCA TGT GTG GCG G) and RdRP_SARSr-R1 (CAR ATG TTA AAS ACA CTA TTA GCA TA). Standard curves were created using plasmid DNA (pEX-A128-RdRP) harboring the corresponding amplicon regions for RdRP target sequence according to GenBank Accession number NC_045512. All quantification experiments have been carried out with biological replicates.

### Neutralization assay

Serum of double mRNA-1273-vaccinated individuals was serially diluted and pre-incubated with 4000 TCID50/mL of SARS-CoV-2 variants at 37°C for 1 h prior transfer to Caco-2-F03 monolayers in 96 well plate. The neutralization titer was determined either by visual scoring of CPE 72 h post infection or caspase 3/7 activity measurement.

### Selection of drug-resistant variant

SARS-CoV-2/FFM3 was serially passaged with increasing concentration (starting concentration - 500nM) of remdesivir in Caco-2-F03. Viral replication was monitored by observation for any cytopathogenic effect present in the culture. Infected cultures were frozen at −80°C and thawed once prior a passaging. Virus was serially passaged by using 1 aliquot of viral stock from the preceding passage to infect fresh Caco-2-F03 cells (MOI of 0.1) in the presence of increasing concentrations of compound for a total of 30 passages, resulting in a strain that could be readily passaged in the presence of remdesivir 2 μM (FFM3^r^REM).

### Sequencing

Extracted nucleic acid was DNase treated, reverse transcribed, and randomly amplified using a Sequence-Independent Single-Primer Amplification (SISPA) method described previously [Lewandowski et al., 2019]. Illumina sequencing used the Nextera XT protocol with 2 × 150-bp paired-end sequencing on a MiSeq.

### Phospholipidosis quantification

Phospholipidosis was assessed as previously described [Tummino et al., 2021]. Cells were treated with hydroxychloroquine in the presence of 7.5 μM nitrobenzoxadiazole-conjugated phosphoethanolamine (NBD-PE) (ThermoFisher). Images were taken and the fluorescence was quantified using a Spark^®^ Mulitmode microplate reader (TECAN).

### Screening assay

The Kinase inhibitor library L-1200 (Selleckchem) containing 1796 compounds was tested in a proof-of-concept screen in Delta-infected Caco-2-F03 cells for the identification of antivirally active agents. Caco-2-F03 cells were seeded into 96-well plates (50,000 cells/well) and incubated at 37°C for 4 days. After the cells reached confluence, the supernatant was replaced by 25 µL/well of medium containing the ABCB1 inhibitor zosuquidar (final concentration 1 µM), 25 µL/well of medium containing kinase inhibitors (final concentration 10 µM) in singlets, and 50 µL/ well SARS-CoV-2 suspension (MOI 0.01). Remdesivir (10 µM) was used as positive control. Plates were incubated at 37°C for 48h prior to the measurement of caspase 3/7 activity as described above. For each plate the Ź score, a measure of statistical effect size and an index for assay quality control, was calculated by: Z′= 1 − (3*s.d.signal + 3*s.d.basal)/(Meansignal − Meanbasal). Only plates with Źscore ≥ 0.5 were further analyzed.

### Drug combination studies

To evaluate antiviral activity of drug combinations, drugs were tested alone or in fixed combinations at 1:2 dilutions using monolayers of Caco-2-F03 cells infected with the indicated SARS-CoV-2 isolates at MOI 1. Antiviral effects were detected 24 h post infection by immunofluorescence staining for S protein. The calculation of IC50, IC75, IC90 and IC95 for single drugs and their combinations as well as combination indices (CIs) was performed using the software CalcuSyn (Biosoft) based on the method of Chou and Talalay [Chou, 2006]. The weighted average CI value (CI_wt_) was calculated according to the formula: CI_wt_ [CI_50_ + 2CI_75_ + 3CI_90_ + 4CI_95_]/10. CI_wt_ values were calculated for mutually exclusive interactions where CI_wt_<1 indicates synergism, CI_wt_ =1 indicates additive effects, and CI_wt_ ˃1 suggest antagonism.

### Statistical analysis

The results are expressed as the mean ± standard deviation of at least three experiments. The Student’s *t*-test was used for comparing two groups. Three and more groups were compared by ANOVA. GraphPad Prism 9 was used to determine IC50 and CC50.

