## Supplementary material for "Identification of novel antiviral drug candidates using an optimized SARS-CoV-2 phenotypic screening platform": Suppl. Figures and Table

Suppl. Figure 1

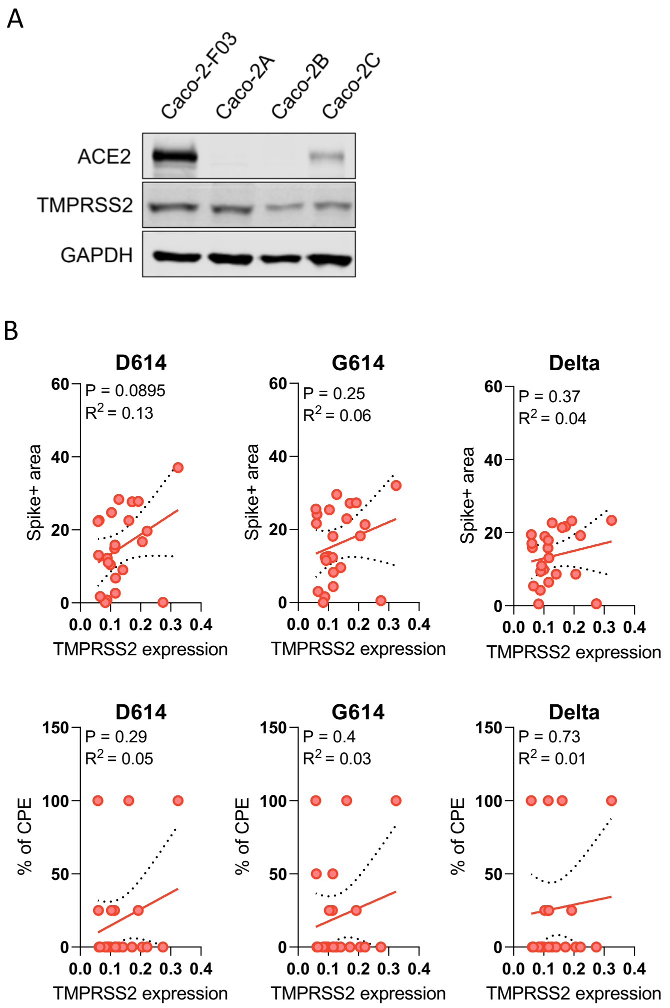


**Suppl. Figure 1. ACE2 and TMPRSS2 levels in Caco-2 cell lines of different origin and correlation of TMPRSS2 levels with SARS-CoV-2 susceptibility of clonal Caco-2A sublines.** A) Western blots indicating ACE2 and TMPRSS2 levels in Caco-2 cell lines of different origin. B) Correlation of TMPRSS2 levels with SARS-CoV-2 susceptibility of clonal Caco-2A sublines as determined by immunostaining for the SARS-CoV-2 S protein and cytopathogenic effect (CPE) formation in SARS-CoV-2/FFM7 (G614) (MOI 1)-infected cells 48h post-infection.

Suppl. Figure 2

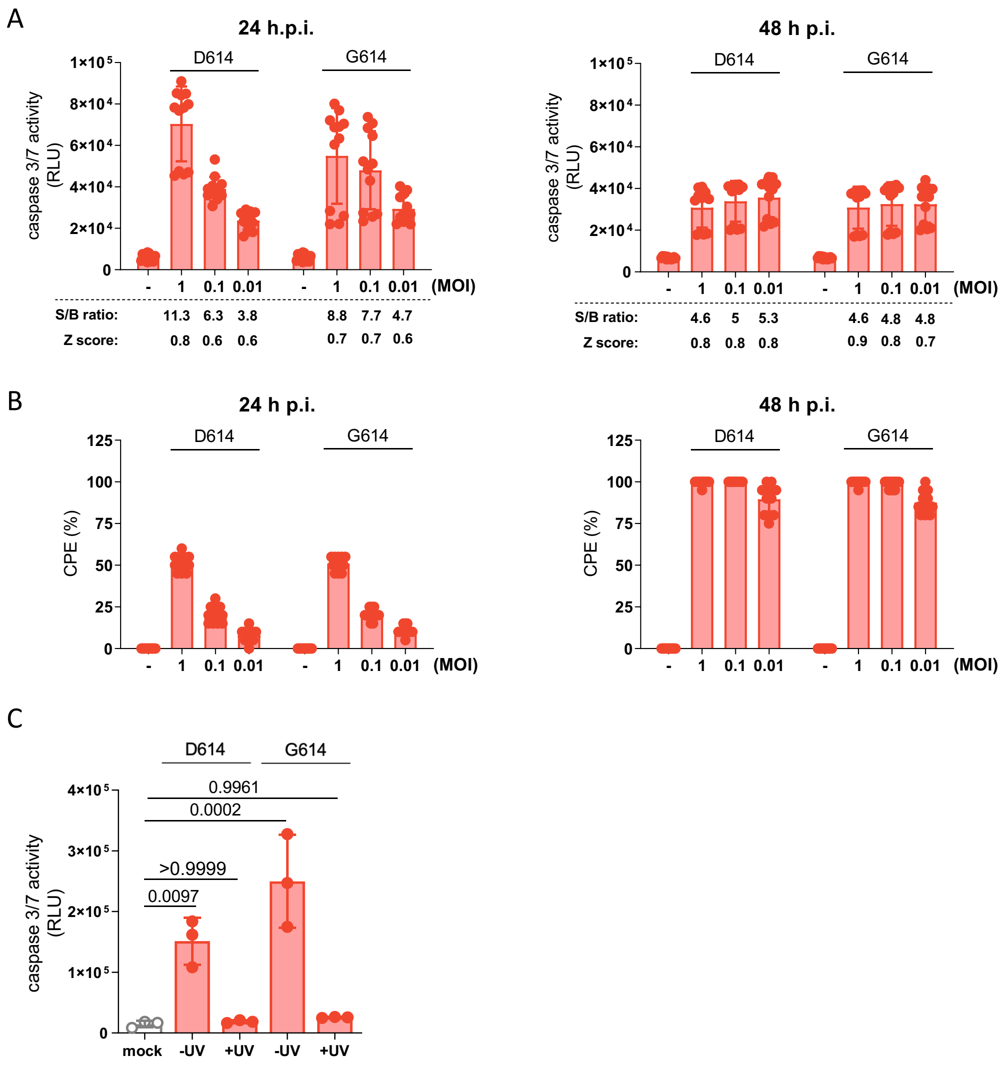


**Suppl. Figure 2. Caspase 3/7 activity as read-out indicating SARS-CoV-2 replication.** A) Caspase 3/7 activity in Caco-2-F03 cells infected with SARS-CoV-2 D614 and G614 isolates at MOI 1, 0.1, and 0.01 as determined by Caspase-Glo assay 24h or 48h post infection, including signal-to-basal (S/B) ratios and Z' scores. B) Cytopathogenic effect (CPE) formation in Caco-2-F03 cells infected with SARS-CoV-2 D614 and G614 isolates at MOI 1, 0.1, and 0.01 24h or 48h post infection. C) Caspase 3/7 activity in Caco-2-F03 cells infected with replication-competent and UV-inactivated SARS-CoV-2 D614 and G614 isolates (MOI 0.01) as determined by Caspase-Glo assay 48h post infection. P-values were calculated by one-way ANOVA.

Suppl. Figure 3

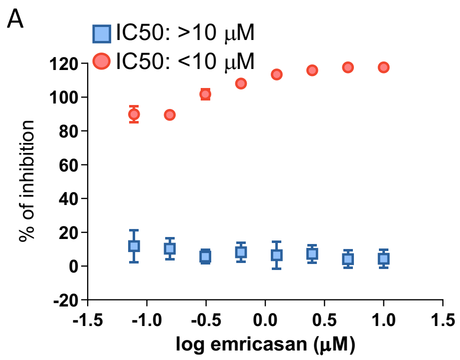


**Suppl. Figure 3. Impact of the pan-caspase inhibitor emricasan on SARS-CoV-2 infection and SARS-CoV-2-mediated caspase 3/7 activity in Caco-2-F03 cells.** Emricasan-induced inhibition of caspase 3/7 activity (red circles) and cellular S protein levels as indicated by immunostaining (blue squares) in G614 (MOI 0.01)-infected Caco-2-F03 cells 48h post infection. Concentrations that reduce caspase 3/7 activity and S staining by 50% (IC50) are also provided.

Suppl. Figure 4

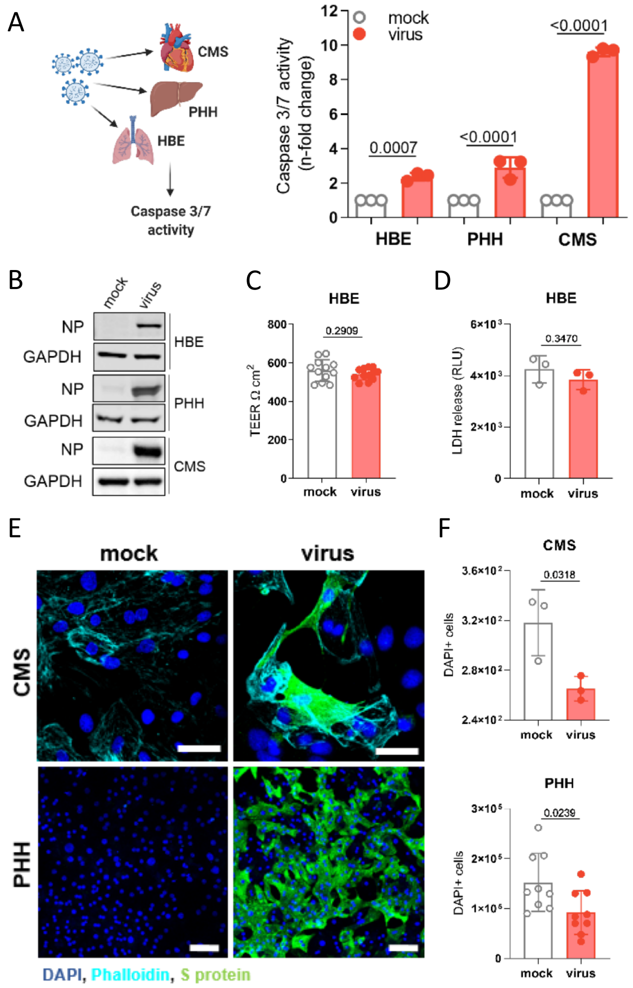


**Suppl. Figure 4. Caspase 3/7 activity in SARS-CoV-2-infected primary human cell cultures**. A) Caspase 3/7 activity in G614 (MOI 1)-infected air liquid interface (ALI) cultures of bronchial epithelial (HBE) cells, cardiomyocytes (CMS) and hepatocytes (PHH) as determined 120h (HBE) or 48h (CMS, PHH) post infection. B) Western blots for the SARS-CoV-2 nucleoprotein (NP) confirming infection in SARS-CoV-2 G614 (MOI 1)-infected primary human cell cultures. C) Transepithelial electrical resistance (TEER) in G614-infected ALI HBE cultures. D) LDH release in G614-infected ALI HBE. C) and D) indicate that SARS-CoV-2 infection does not result in a CPE in ALI HBE cultures. E) CPE formation in G614-infected CMS and PHH. Immunofluorescence staining indicates SARS-CoV-2-infected cells by S protein levels and cells by DAPI and phalloidin staining. F) Quantification of DAPI-stained nuclei in G614-infected CMS and PHH. All p values were calculated by unpaired t-test.

Suppl. Figure 5

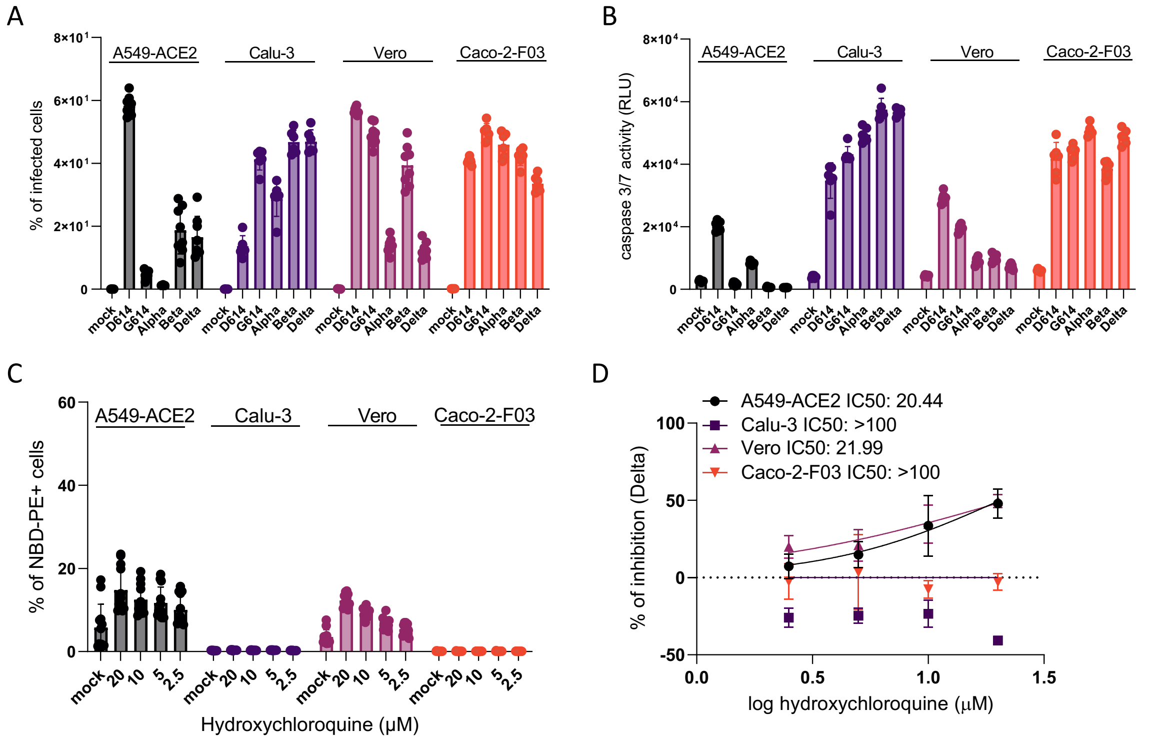


**Suppl. Figure 5. Susceptibility to different SARS-CoV-2 variants and drug-induced phospholipidosis in different cell lines used for SARS-CoV-2 cultivation.** A) Spike (S) protein levels as determined by immunostaining and B) caspase 3/7 activity in cell lines infected with different SARS-CoV-2 isolates at MOI 0.01 at 48h post infection. C) Hydroxychloroquine-induced phospholipidosis as indicated by nitrobenzoxadiazole-conjugated phosphoethanolamine (NBD-PE) staining. D) Effects of hydroxychloroquine on cellular S levels in SARS-CoV-2 Delta (MOI 0.01)-infected cells 48h post infection.

Suppl. Figure 6

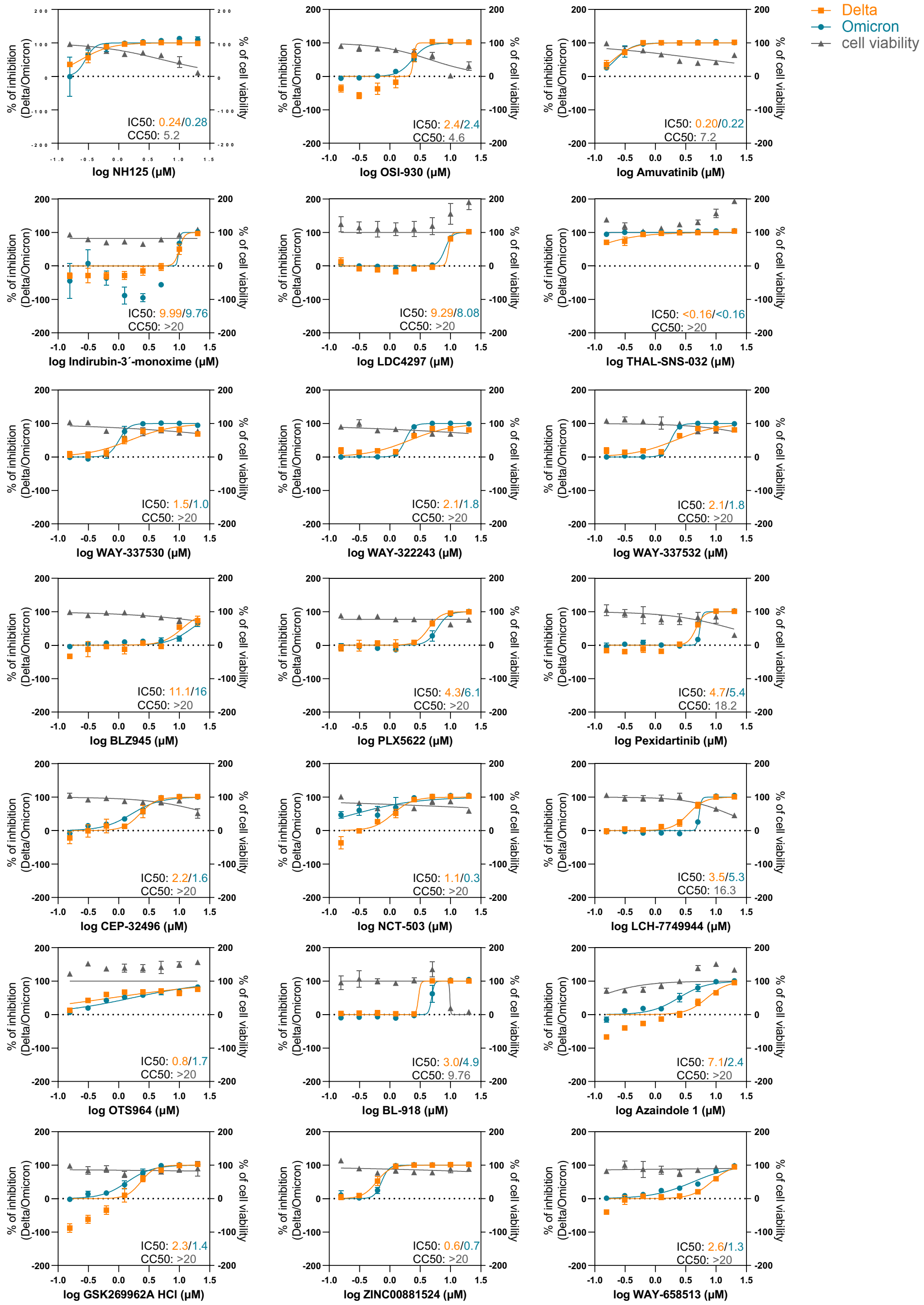


**Suppl. Figure 6. Dose-response curves confirming the anti-SARS-CoV-2 activity of 21 hits by the determination of drug-response curves in in SARS-CoV-2 strain FFM3 (MOI 0.01)-infected Caco-2-F03 cells using immunostaining for the viral S protein as read-out 48h post infection.** Cell viability was determined by CellTiterGlo assay.

Suppl. Figure 7

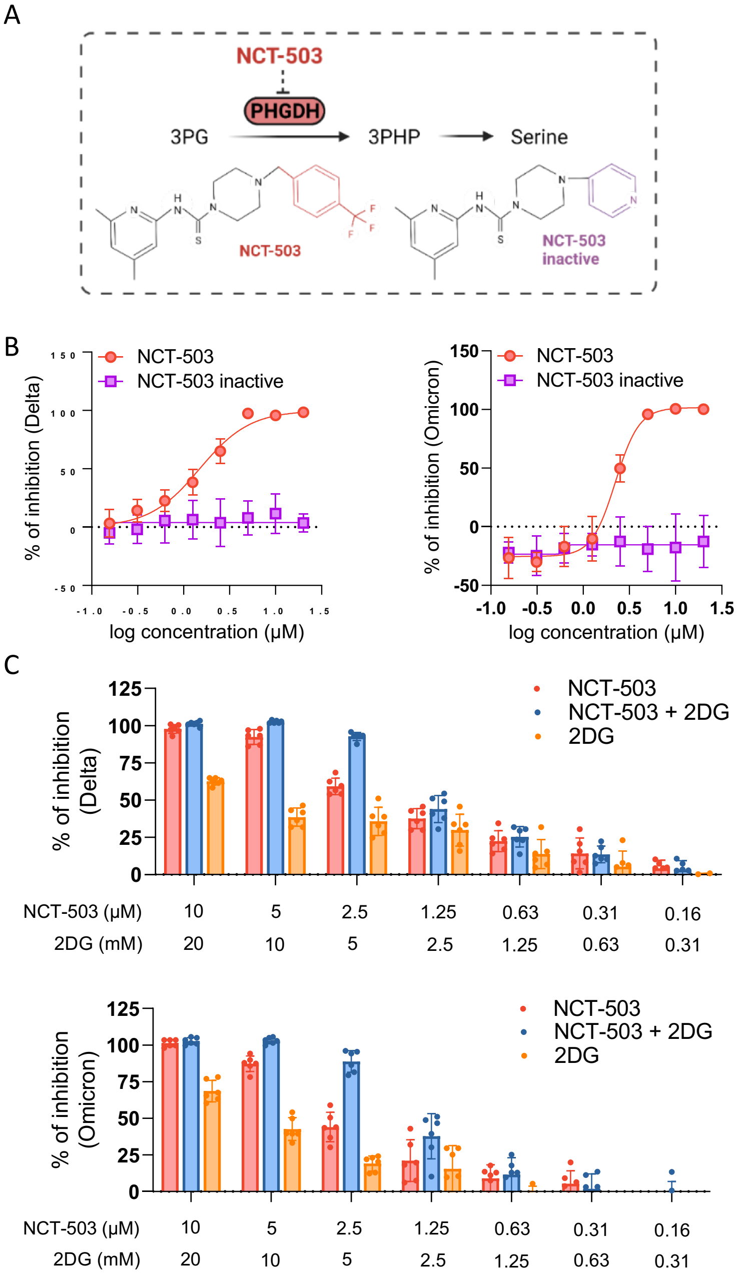


**Suppl. Figure 7. Investigation of the anti-SARS-CoV-2 effects of the PHGDH inhibitor NCT-503.** A) Chemical structure of NCT-503 and a chemically closely related control that does not inhibit PHGDH. B) Dose-response curves indicating the anti-SARS-CoV-2 activity of NCT-503 and the inactive control in Caco-2-F03 cells infected with a Delta and an Omicron isolate (MOI 0.01) as determined by immunostaining for S 48h post infection. C) Combined antiviral effects of NCT503 and 2DG in Delta and

Omicron (MOI 0.01)-infected Caco-2-F03 cells as determined by S immunostaining 48h post infection.

**Suppl. Table 1. Short tandem repeat profiles of Caco-2 cell lines from different sources.**

| | TH 01 | D5 S818 | D13 S317 | D7 S820 | D16 S539 | CSF1 PO | Amel | VWA | TPOX |
|---|---|---|---|---|---|---|---|---|---|
| Reference profile | 6; 6 | 12; 13 | 11,13; 14 | 11; 12 | 12; 13 | 11; 11 | X; X | 16; 18 | 9; 11 |
| Caco-2-F03 | 6; 6 | 12; (13) | (11,13); 14 | 11; 12 | 12; 13 | 11; 11 | X; X | 16; 18 | 9; 11 |
| Caco-2A | 6; 6 | 12; (13) | 11,13; 14 | 11; 12 | 12; 13 | 11; 11 | X; X | 16; 18 | 9; 11 |
| Caco-2B | 6; 6 | 12; 13 | 11,13; 14 | 11; 12 | 12; 13 | 11; 11 | X; X | 16; 18 | 9; 11 |
| Caco-2C | 6; 6 | 12; (13) | 11,(13); 14 | 11; 12 | 12; 13 | 11; 11 | X; X | 16; 18 | 9; 11 |
